# Age-dependent dysregulated post-ovulatory debris clearance promotes ovarian fibroinflammation

**DOI:** 10.64898/2026.09.23.753904

**Authors:** Aubrey Converse, Madeline J. Perry, Shweta S. Dipali, Prianka H. Hashim, Jennifer L. Gerton, Michele T. Pritchard, Francesca E. Duncan

## Abstract

The ovary is one of the first organs to age, with declines in oocyte quantity and quality leading to infertility and loss of endocrine function. Why the ovary ages earlier than somatic tissues has remained enigmatic but may relate to its highly dynamic physiology and substantial demand for cellular debris clearance. Each ovulation forms a corpus luteum (CL), a transient endocrine structure which supports pregnancy but regresses in its absence. Because CLs have a short functional lifespan but prolonged structural regression, they represent a substantial source of ovarian debris. We hypothesized that post-ovulatory debris clearance becomes impaired with age and contributes to ovarian fibroinflammation. Using a CL-labeling system, we demonstrated that CLs from reproductively old mice show dysregulated regression, resulting in an accumulation of persistent CLs (pCLs) which appeared concurrently with ovarian multinucleated giant cells (MNGCs). Laser capture microdissection, RNA sequencing, and histological evaluation of CLs, pCLs, and MNGCs demonstrated that pCLs were focal hotspots of proinflammatory and debris-associated signatures enriched in lipids, extracellular matrix, and immune cells. Although pCLs retained luteal features, they were transcriptionally more similar to MNGCs than CLs. Consistent with this, an *ex vivo* CL model acquired MNGC-associated molecular features during extended culture, suggesting that pCLs may represent sites of MNGC formation. Culture of isolated follicles or ovarian stromal organoids with factors from MNGC-enriched explants induced compartment-specific fibroinflammatory responses. These findings provide novel mechanistic insight into ovarian aging, whereby post-ovulatory debris clearance is compromised with age, leading to pCL accumulation, MNGC formation, and a fibroinflammatory milieu.

## INTRODUCTION

The ovary is one of the first organs to exhibit an age-dependent functional decline. Ovarian aging begins in women in their mid-thirties, and reproductive function ceases completely at menopause. Ovarian aging hallmarks include reduced oocyte quantity and quality, decreased endocrine output, increased stromal fibrosis, and development of a proinflammatory milieu (1–6). Together, these age-associated fibrotic and inflammatory changes, or “fibroinflammaging,” can directly impact follicle growth and development, gamete quality, and ovulation (7–9). Why the ovary ages prior to somatic tissues has remained enigmatic, but the origins of this phenomenon likely lie in the ovary’s extraordinarily dynamic nature. Human females are born with a finite pool of millions of follicles, but only a few hundred will reach a terminal stage and be ovulated, with the majority ultimately being eliminated by cell death mechanisms (10, 11). Nevertheless, women will still ovulate every 28 days over their reproductive lifespans, resulting in a total of approximately 400 ovulatory events and repetitive cycles of damage and wound healing (12). Following ovulation, the residual follicle wall involutes and undergoes a process known as luteinization to form the corpus luteum (CL), a transient endocrine structure which produces progesterone in preparation for potential pregnancy (13, 14). If pregnancy does not occur, the CL ceases its endocrine activity and undergoes luteolysis or structural regression. Together, follicular atresia, ovulation and post-ovulatory wound repair, and CL regression impose a substantial and recurrent debris burden on the ovary.

Tissue homeostasis relies on the coordinated turnover of macromolecules, organelles, dying cells, and cellular debris (15, 16). With age, there is an inability to maintain effective tissue homeostasis and this contributes to cellular senescence, loss of proteostasis, mitochondrial dysfunction, and chronic inflammation (17–20). Defective debris clearance is also a feature of age-related conditions, including Alzheimer’s disease, Parkinson’s disease, atherosclerosis, arthritis, and cancer (21–26). Given the ovary’s exceptionally high demand for tissue remodeling and debris clearance, age-associated defects in these homeostatic processes may underly and contribute to this organ’s accelerated aging and development of fibroinflammatory phenotypes. CLs are likely major contributors to ovarian cellular debris burden because they have a limited functional lifespan but prolonged structural regression due to their large size. In mice, loss of steroidogenic activity, or functional CL regression, occurs within 3 days post-ovulation, whereas complete structural CL regression, which requires coordinated vascular remodeling, immune-cell recruitment, cell death, and phagocytic clearance, can take up to three estrous cycles (13, 27–31). In humans, the corpus albicans or scar-like tissue remaining from the degenerating CL can persist for months (32). However, whether CL regression and post-ovulatory debris clearance are affected with age remains unknown. This is in part due to the difficulty of studying structural CL regression due to the extended non-functional lifespan of these structures and the inability to decipher CLs from different ovulatory cycles.

In the current study, we address the hypothesis that post-ovulatory debris clearance becomes impaired with age and contributes to ovarian fibroinflammation. To carry this out, we developed an innovative strategy to track CLs in mice over multiple estrous cycles and demonstrated that CL regression is dysregulated with age, resulting in the accumulation of persistent CLs (pCLs). As pCLs occur concurrently with age-associated multinucleated giant cells (MNGCs), we sought to determine the relationship between CLs, pCLs, and MNGCs by utilizing laser capture microdissection and bulk RNA Sequencing (RNASeq). While pCLs expressed a unique profile of genes involved in efferocytosis, lipid efflux, and extracellular matrix (ECM) organization, they also expressed canonical CL and MNGC markers. Principal component analysis revealed that pCLs were transcriptomically most similar to MNGCs despite their CL lineage and may provide sites that promote MNGC formation. This was further supported by an ex vivo CL model which also acquired MNGC-associated features. Exposure of *ex vivo* models of distinct ovarian subcompartments to factors released by MNGC-enriched explants resulted in the upregulation of genes involved in inflammation in follicles and tissue and ECM remodeling in the stroma, recapitulating what is observed in ovarian aging. In addition, long-term follicle culture in the presence of MNGC-secreted factors resulted in altered growth and impaired oocyte maturation relative to controls. These findings define dysregulated post-ovulatory debris clearance as a novel mechanism underlying age-dependent MNGC formation and ovarian fibroinflammation and lay the foundation for novel therapeutics targeting debris clearance to enhance ovarian longevity.

## RESULTS

### CL regression is impaired with advanced reproductive age

While functional CL regression is well defined, the kinetics of structural regression have been more difficult to assess (14, 27, 31). In the mouse, CLs can take up to three estrous cycles to fully regress (27). Given that mice are polyovulatory, the ovary contains CLs from numerous ovulatory events at any given time, posing a challenge in determining the post-ovulatory age of specific CLs. Therefore, we developed a robust tracking strategy that leveraged the unique proliferative profile of the CL during its formation (Fig. 1A). CLs undergo high levels of angiogenesis and luteal cell proliferation at the time of ovulation, after which little to no mitotic activity is observed (33–37). Therefore, we injected bromodeoxyuridine (BrdU), a thymidine analog that incorporates into DNA during S-phase, into mice following a hormonally induced ovulatory event to tag newly formed CLs. With this strategy, the BrdU label is only incorporated into CLs formed from the induced ovulatory event, as older CLs are no longer proliferative. Furthermore, the BrdU label can be retained for extended periods due to the lack of mitotic dilution after initial CL formation (Fig. 1A). To determine the effect of age on CL regression kinetics, we performed BrdU injections in reproductively young and old mice following superovulation. Ovaries were then analyzed from mice at 1 day and 1, 2, and 3 wks post-ovulation. Of note, while positive BrdU-labeling was observed in follicles at all stages of development 1 day post-ovulation, no follicular labeling was observed at later timepoints, indicating that BrdU is rapidly diluted in proliferating granulosa cells (Fig. 1B). Therefore the BrdU-label in follicles does not contribute to positive signal in CLs formed in subsequent ovulatory events (Fig. 1B). BrdU-labeled CLs were observed in ovaries at all timepoints, including at 3 wks consistent with the extended timecourse of CL regression (Fig. 1C). To determine the kinetics of CL structural regression, the number of BrdU-positive CLs was counted in each ovary across age and timepoints. In reproductively young mice, the number of labeled CLs decreased over time, with a 73% decrease in the number of BrdU-positive CLs at 3 wks post-ovulation compared to the 1 day timepoint (Fig. 1D). With advanced age, there were fewer BrdU-positive CLs at 1 day post-ovulation relative to young counterparts, which was expected given the age-dependent decline in follicle numbers (Fig. 1E) (8). However, the number of BrdU-positive CLs remained constant across the 3 wk timeframe in reproductively old mice, indicative of a delay or dysregulation of CL regression and clearance (Fig. 1E).

**Figure 1.**
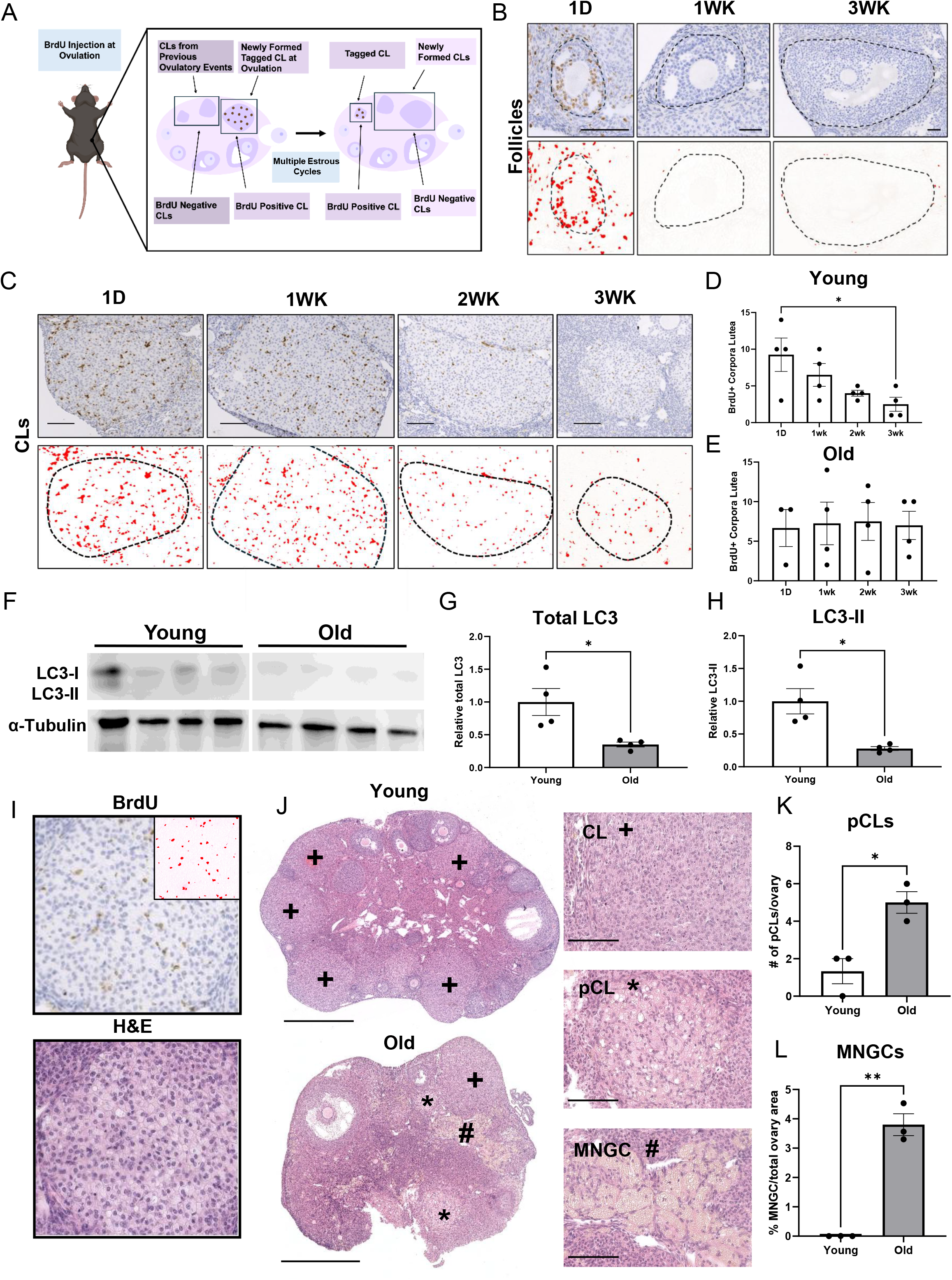
Dysregulated corpora lutea regression contributes to post-ovulatory debris accumulation in the aged ovary. (A) Schematic of BrdU-based corpora lutea (CL) tagging system. (B) Representative images of BrdU-labeled follicles at 1 day, 1 wk, and 3 wks post-ovulation. (C) Immunohistochemistry of CL BrdU labeling across all timepoints in old mouse ovaries and corresponding thresholded images showing positive stained area (red). Quantification of BrdU-labeled CLs in (D) young and (E) old mouse ovaries at 1 day, 1 wk, 2 wks, and 3 wks post-ovulation. Western blot (F) and quantification of total LC3 (G) and active LC3-II (H) in 1 wk post-ovulatory CLs normalized to α-Tubulin. (I) Representative image of a BrdU-labeled CL at 3 wks post-ovulation and corresponding morphology as observed by H&E staining. Inset represents thresholded BrdU signal. (J) Representative ovarian sections from young and old mice indicating CLs (+), persistent CLs (pCL,*), and multinucleated giant cells (MNGC,#). Quantification of pCL number (K) and MNGC area (L) in young and old ovarian sections. Data represent means ± SEM, with 3-4 biological samples assessed for all analyses. For (D) and (E), data were tested for normality using the Shapiro-Wilk test, then analyzed by one-way ANOVA with Tukey’s multiple comparison post-test (normal) or by Kruskal-Wallis test with Dunn’s multiple comparisons post-test (nonparametric). For (G), (H), (K), and (L), significance was determined by Welch’s *t* test. (*, *P* < 0.05; **, *P* < 0.01). Scale bars: 100 µm for (B) and (C); 500 µm for whole ovarian sections (J) and 100 µm for insets.

Tissues utilize various mechanisms to ensure proper debris clearance. Autophagy is the process by which cells degrade cellular components through a lysosomal-dependent mechanism to remove and recycle damaged and dysfunctional proteins, organelles, and other molecular components (22, 38). Autophagy plays a prominent role in CL regression (39, 40). To determine whether defects in autophagy may contribute to dysregulated post-ovulatory debris clearance with age, we examined the expression of LC3, an essential autophagosome membrane component, in microdissected CLs from reproductively young and old mice. Western blot analyses revealed an age-associated decline in expression of both total-LC3 and the active form LC3-II in CLs (Fig. 1F-H) (41). These findings are consistent with reduced autophagy-associated machinery and activity, which likely contribute to the age-associated dysregulation of CL regression.

### Aging is associated with persistent CL and multinucleated giant cell accumulation

At the 2 wk and 3 wk timepoints, the BrdU-positive CLs exhibited a high cytoplasmic-to-nuclear ratio, spherical shape, and vacuolar or foamy appearance (Fig. 1I), a morphology that has been reported for late-stage regressing CLs (32, 42). Consistent with the age-associated dysregulation in CL regression, we observed a 3.8-fold enrichment of CLs with this particular appearance, which we termed persistent CLs (pCLs), in ovaries from reproductively old mice relative to young counterparts (Fig. 1J,K). These data provide further evidence that CL regression becomes dysregulated with advanced reproductive age and results in the accumulation of pCLs or age-associated ovulatory debris burden. Interestingly, the enrichment of pCLs in ovaries from reproductively old mice occurred concurrently with ovarian multinucleated giant cells (MNGCs) (Fig. 1J,L), which arise from macrophage fusion events and are implicated in debris clearance and chronic inflammation in ovarian aging, autoimmune diseases, at sites of infection, and in response to foreign bodies (43–45). Ovarian MNGCs are age-associated, form complexes with T cells, and are characterized by a proinflammatory and degradation-associated gene expression signature (45). Thus, the simultaneous presence of this unique degradation-associated cell population with pCLs suggests that MNGCs may arise in the aging ovary to compensate for the loss of homeostatic debris clearance mechanisms.

### pCLs are molecularly distinct, lipid-rich, fibrotic, inflammatory structures

To investigate the molecular relationship between pCLs, MNGCs, and CLs, we utilized laser capture microdissection (LCM) to isolate these specific structures from ovarian tissue sections from reproductively old mice and then performed transcriptomic analyses using bulk RNASeq (Fig. 2A). Images of H&E-stained serial sections were also used to confirm structure identification prior to capture (Supplemental Fig. 1i-iii). In unstained tissue sections, CLs had a darker appearance and exhibit well-defined borders (Supplemental Fig. 1iv). pCLs did not have a defined border and were lighter in appearance with dark vacuolar spaces (Supplemental Fig. 1v), while MNGCs were darkly pigmented and irregularly shaped (Supplemental Fig. 1vi).

**Figure 2.**
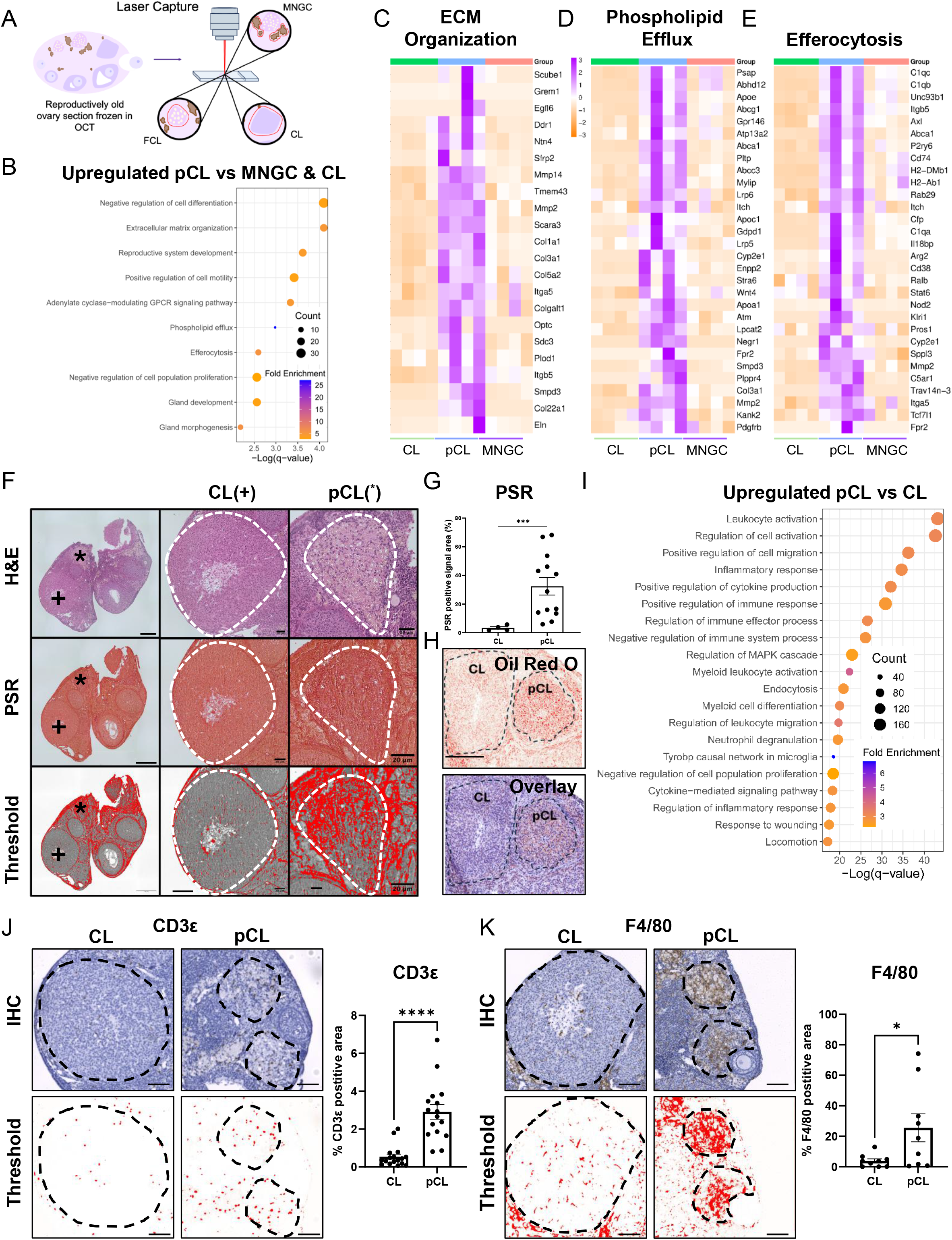
Persistent CLs are molecularly distinct debris-rich fibroinflammatory structures. (A) Schematic of laser capture microdissection workflow. (B) Pathway analysis of upregulated DEGs in pCLs compared to both CLs and MNGCs. Heatmaps of DEGs (pCL vs CL/MNGC) pertaining to extracellular matrix (ECM) organization (C), phospholipid efflux (D), and efferocytosis (E). (F) Representative images and (G) quantification of picrosirius red staining (PSR) of CLs (+) and pCLs (*) in old ovaries. (H) Representative image of Oil red O staining of CLs and pCLs from old ovaries. (I) Pathway analysis of upregulated DEGs in pCLs compared to CLs. Immunohistochemical analysis and quantification of T cells (CD3ε) (J) and macrophage (F4/80) marker protein expression in CLs and pCLs from old mice. Bottom panels show positive area (red) after deconvolution and thresholding. Data represent means ± SEM, with 3-4 biological samples assessed for all analyses. For (G), (J), and (K), a minimum of 1 and a maximum of 5 CLs or pCLs were quantified for ≥ 3 biological ovarian samples. Significance was determined by Welch’s *t* test. (*, *P* < 0.05; ***, *P* < 0.001; ****, *P* < 0.0001). Scale bars: (F) 100 µm for whole ovarian section and 20 µm for higher magnification CL and pCL images; (H) 100 µm; (J-K) 100 µm.

Transcriptomic profiling indicated that pCLs showed upregulation of 372 genes compared to MNGC and CL samples (p<0.05; log_2_ fold change ≥ 0.5). Unbiased pathway analysis revealed an enrichment of processes consistent with debris burden in pCLs compared to both CLs and MNGCs, including “extracellular matrix organization,” phospholipid efflux,” and “efferocytosis” (Fig. 2B). DEGs contributing to these pathways were visualized by heatmaps (Fig. 2C-E). These gene signatures indicate that pCLs may be sites of dysregulated extracellular matrix remodeling and lipid handling, with enhanced debris clearance signaling. To validate these molecular signatures, collagen I and III and neutral lipid droplets were visualized in ovarian tissue sections using Picrosirius red (PSR) and Oil Red O staining. pCLs were enriched for collagen I and III compared to CLs (Fig. 2F,G), indicating that these sites represent highly fibrotic foci and likely contribute to the age-associated increases in ovarian fibrosis and stiffness that have been previously reported (1, 2). Additionally, pCLs contained lipid droplets that appeared larger and more densely packed than those in CLs (Fig. 2H), suggesting that lipid metabolic and trafficking processes are dysregulated at these sites. Due to these prominent differences in intracellular and extracellular debris burden between pCLs and CLs, we next assessed how pCLs and CLs differ transcriptionally. pCLs and CLs exhibited 3,302 DEGs with an adjusted *P*-value < 0.05 and log_2_ fold change ≥ 1 (2,241 upregulated and 1,061 downregulated in pCLs compared to CLs; Supplemental Fig. 2A). Pathway analysis of DEGs upregulated in pCLs compared to CLs indicated various immune-related processes, including “leukocyte activation,” “positive regulation of cytokine production,” and “inflammatory response,” indicating that pCLs have increased immune and proinflammatory signatures compared to CLs (Fig. 2I). To determine whether these gene signatures reflect increased immune cell presence within pCLs, T cells and macrophages were assessed through immunolocalization. T cell (CD3ε) and macrophage (F4/80) populations were significantly increased in pCLs compared to CLs, indicating enhanced immune cell recruitment and/or retention to these structures (Fig. 2J,K). Overall, the distinct transcriptomic profile, and molecular and cellular composition of pCLs indicate they represent fibroinflammatory foci within the aging ovarian microenvironment, characterized by immune cell accumulation and persistent cellular and extracellular debris.

### pCLs are transcriptionally similar to MNGCs

While over 3,000 DEGs were found between pCLs and CLs, only 991 genes were differentially expressed between pCLs and MNGCs (717 upregulated and 274 downregulated in pCLs compared to MNGCs; Supplemental Fig. 2B). Principal component analysis (PCA) revealed distinct clustering of CLs, pCLs, and MNGCs (Fig. 3A). However, despite their luteal origin, pCLs clustered closer to MNGCs than CLs, indicating greater transcriptomic similarity between pCLs and MNGCs (Fig. 3A). Consistent with this observation, global transcriptome correlation analyses demonstrated strong similarity between pCLs and MNGCs, with correlation coefficients between 0.59 and 0.78, whereas correlation coefficients between CL and pCL groups were between 0.06 and 0.43. Of note, several pCL-MNGC comparisons exhibited correlation coefficients comparable to those observed between MNGC samples (0.72-0.96) (Fig. 3B). To further examine the transcriptomic relationship among CLs, pCLs, and MNGCs, we focused on genes associated with lipid metabolic processes because this was the top GO term of genes upregulated in CLs relative to pCLs as well as immune system processes which is a broader functional category that encompassed the immune-related signatures exhibited by pCLs. Consistent with the PCA and correlation analyses, pCLs exhibited functional gene expression patterns that were more like those of MNGCs relative to CLs, characterized by reduced expression of genes involved in lipid metabolism and increased expression of immune-associated genes (Fig. 3C,D). Overall, these findings demonstrate that pCLs are transcriptionally distinct from CLs and share several immune-related features with MNGCs, supporting a close association between these structures within the aging ovary.

**Figure 3.**
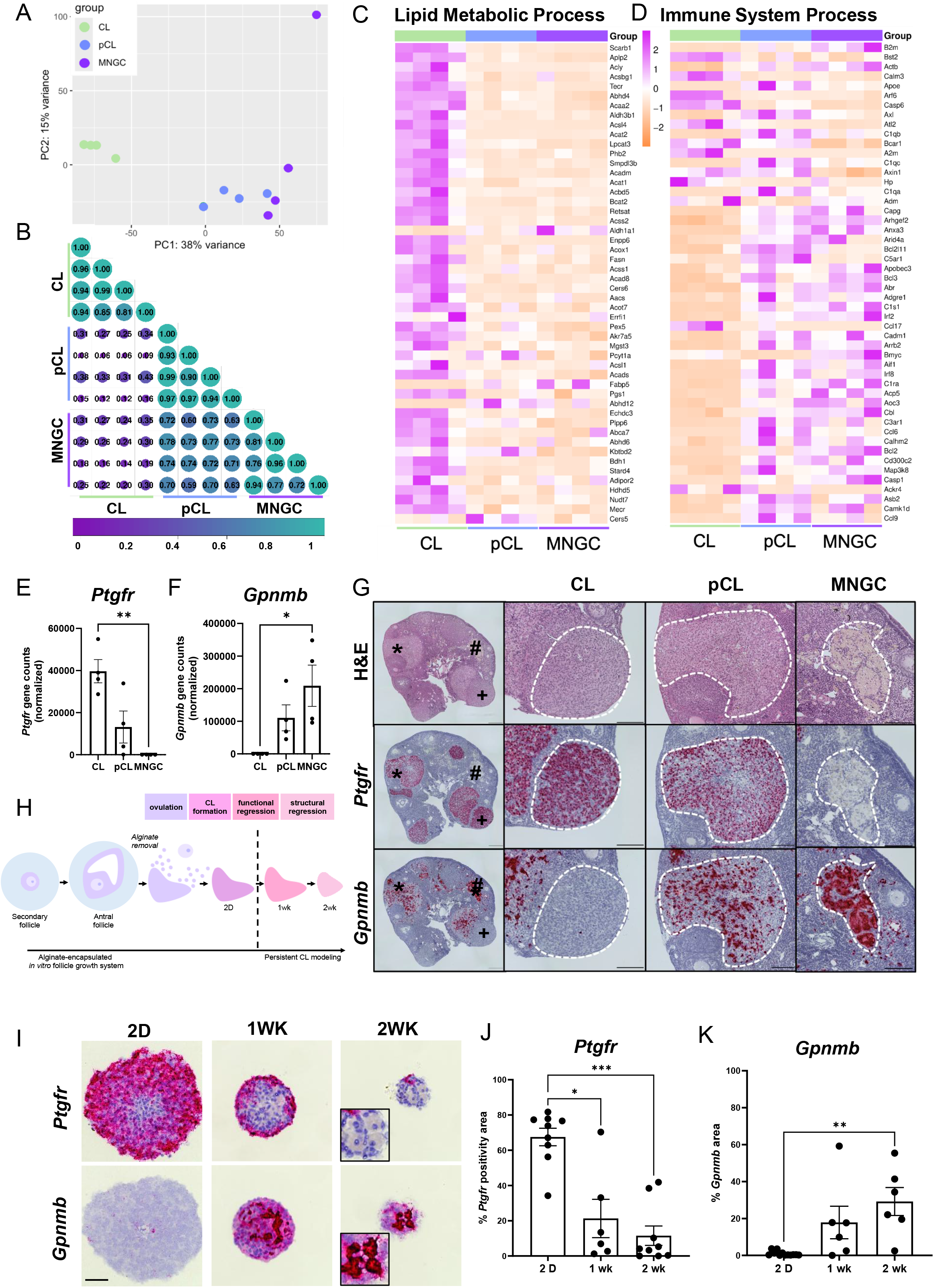
pCLs are molecular intermediaries between CLs and MNGCs. (A) Principal component analysis visualizing variance between CLs (green), pCLs (blue), and MNGCs (purple). (B) Correlation analysis comparing CLs, pCLs, and MNGCs. Heatmaps of DEGs (pCL vs CL) pertaining to lipid metabolic processes (C) and immune system processes (D). RNAseq gene counts for *Ptgfr* (E) and *Gpnmb* (F). (G) Representative RNAscope visualizing transcripts of *Ptgfr* and *Gpnmb* in CLs (+), pCLs (*), and MNGCs (#) within aged ovarian tissue sections. (H) Schematic of the *ex vivo*-derived persistent CL model. (I) Representative RNAscope visualizing transcripts of *Ptgfr* and *Gpnmb* in *ex vivo*-derived CLs at 2 day, 1 wk and 2 wk post-ovulation. Insets indicated pigmented *Gpnmb*-positive cells with putative multinucleation. Quantification of *Ptgfr* (J) and *Gpnmb* (K) in *ex vivo-*derived CLs across the post-ovulatory time course. Data represent means ± SEM, with 3-4 biological samples assessed for all analyses. For (E), (F), (J), and (K) data were tested for normality using the Shapiro-Wilk test, then analyzed by one-way ANOVA with Tukey’s multiple comparison post-test (normal) or by Kruskal-Wallis test with Dunn’s multiple comparisons post-test (nonparametric). For (J) and (K), 2-4 *ex vivo*-derived CLs were quantified for 3 biological samples. (*, *P* < 0.05; **, *P* < 0.01; ***, *P* < 0.001). Scale bars: (G) and (I) 100 µm for all panels.

Although pCLs are derived from CLs, their transcriptomic profiles exhibit greater similarity to those of MNGCs. We therefore examined their expression of luteal- and MNGC-specific markers. *Ptgfr* encodes the prostaglandin F receptor, which is essential for luteolysis and only expressed by luteal cells within the ovary (39, 40, 46–48), whereas *Gpnmb* (glycoprotein NMB) is a marker of ovarian and non-ovarian MNGCs (45, 49, 50). Consistent with their distinct cellular identities, bulk RNAseq data demonstrated high *Ptgfr* expression in CLs and no detectable expression in MNGCs, while *Gpnmb* exhibited an opposite expression pattern, with high expression in MNGCs but no detectable expression in CLs (Fig. 3E,F). Notably, pCLs displayed intermediate expression levels of both genes, retaining expression of the luteal-specific marker *Ptgfr*, while simultaneously expressing the MNGC marker *Gpnmb*. *In situ* hybridization confirmed these expression and localization patterns, with robust expression of *Ptgfr* in CLs and pCLs, whereas *Gpnmb* was present in MNGCs and pCLs but not in CLs (Fig. 3G). These data indicate that pCLs share features with both CLs and MNGCs, supporting a close molecular relationship and identifying pCLs as potential sites of MNGC formation in the aging ovary.

To determine whether MNGC-associated characteristics emerge during CL persistence, we next utilized an *ex vivo* CL regression system to model CL persistence. This model utilizes encapsulated *in vitro* follicle growth (eIVFG) to generate antral follicles that can be induced to ovulate, luteinize, and form functionally active CLs (51–53). Because this *ex vivo* system does not recapitulate the intact vascular, stromal, and recruited immune-cell compartments that coordinate structural luteolysis *in vivo* (29, 30), structural regression remains incomplete, allowing the development of a persistent CL phenotype by 1 wk post-ovulation (Fig. 3H). *In situ* hybridization of luteal- and MNGC-specific markers revealed that *ex vivo*-derived CLs recapitulated the expression patterns observed in pCLs *in vivo*. *Ex vivo*-derived CLs exhibited high *Ptgfr* expression at 2 days post-ovulation, consistent with the initiation of functional regression, and a steady and sustained decrease in expression at 1 and 2 wks post-ovulation (Fig. 3I,J). Interestingly, little to no *Gpnmb* expression was observed at 2 days post-ovulation, whereas expression was apparent in most CLs by 1 wk and prominent by 2 wks post-ovulation (Fig. 3I,K). Additionally, *Gpnmb*-positive cells that also had a pigmented appearance and potential multinucleation were observed at 2 wks post-ovulation (Fig. 3I). These findings demonstrate that MNGC-associated molecular features emerge in persistent *ex vivo* CL structures, supporting their identification as putative sites of MNGC formation.

### Secreted factors from MNGC-enriched explants drive fibroinflammatory changes to stromal and follicular compartments

We previously demonstrated that ovarian MNGCs are highly prevalent within the aging ovary and possess a transcriptomic profile distinct from other ovarian macrophages, characterized by enhanced debris-degradative and immune-signaling functions (45). Our current findings suggest that MNGCs arise at sites of persistent ovulatory debris, consistent with their unique degradation-related signatures. Given their penetrance and abundance within the aging ovary, we next sought to define the functional impact of MNGCs on the surrounding ovarian microenvironment. To address this, we established an *ex vivo* MNGC-enriched explant culture system and examined the effects of factors secreted by these explants on models of the ovarian follicular and stromal compartments (54, 55) (Fig. 4A).

**Figure 4.**
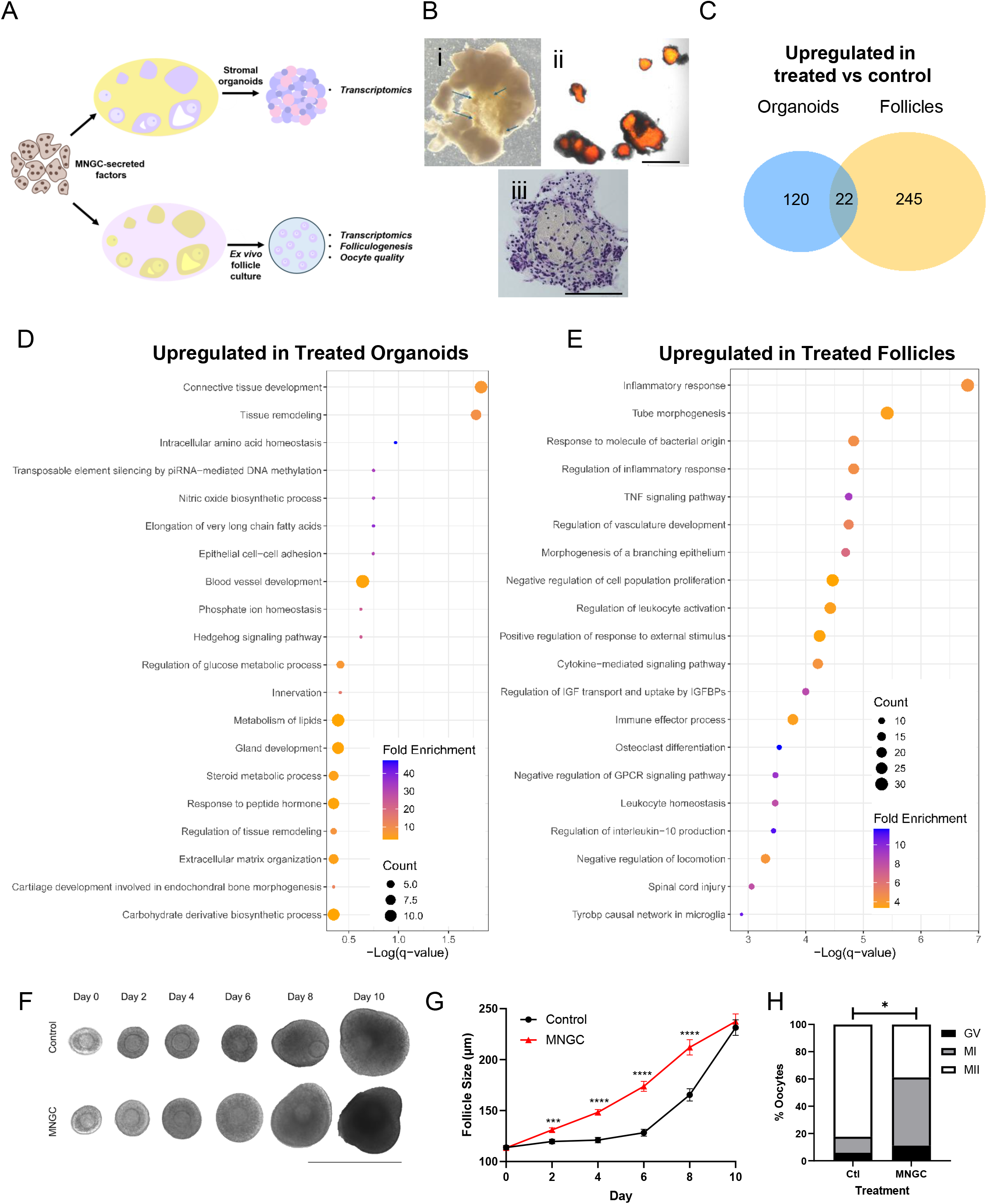
MNGC-enriched explant secreted factors mediate fibroinflammation of stromal and follicle compartments. (A) Schematic of MNGC-enriched explant secreted factor effect on *in vitro* models of ovarian stromal and follicular subcompartments. (B) Brightfield image of aged ovary indicating pigmented putative MNGCs indicated by blue arrows (i), autofluorescence of MNGC-enriched fractions (ii), confirmation of MNGC enrichment post-fixation (iii). (C) Venn diagram comparing DEGs that were upregulated with treatment with conditioned medium compared to control in follicle and stromal compartments. Pathway analysis of DEGs that were upregulated in organoids (D) and follicles (E) treated with MNGC-enriched explant conditioned medium compared to controls. Representative images (F) and quantification (G) of follicle-growth in the presence or absence of MNGC-enriched explant conditioned medium. (H) Meiotic staging of oocytes obtained after maturation of follicles cultured in the presence or absence of MNGC-enriched explant conditioned medium. MNGC: Treatment with MNGC-enriched explant conditioned medium; GV: germinal vesicle; MI: metaphase I-arrest; MII: metaphase II-arrest. All data are presented as mean ± SEM or the proportion of the total oocytes assessed. Two-tailed Student’s t-test was used to analyze (G) while a χ2 test was used to analyze (H) (*,*P* <0.05; ***, *P* <0.001, ****, *P* <0.0001). For (C-E) the experiment was repeated 4 times. For (G-H), each treatment was repeated 3-4 times with 10 follicles each. For (H) 17-18 oocytes were assessed across 3-4 biological trials. Scale bars: (Bii) 500 µm; (Biii) 100 µm; (F) 400 µm.

MNGCs were identified in aged ovaries under brightfield microscopy by their pigmented appearance, which is due to their high lipofuscin content (6). MNGC-enriched fractions were microdissected and their identity was confirmed by their characteristic autofluorescence (Fig. 4Bi-ii) (45). In addition, histological examination further validated MNGC enrichment in these tissue explants (Fig. 4Bii-iii). As these MNGC-enriched fractions also likely contain MNGC-associated T cells and other stromal cell types, we termed these fractions MNGC-enriched explants (45). We first performed targeted cytokine profiling of explant-conditioned medium, which demonstrated the presence of several proinflammatory cytokines including IL-6, CXCL1, and CCL2 (Supplemental Fig. 3). We then used conditioned medium from MNGC-enriched explants to treat ovarian stromal organoids and isolated secondary follicles to directly determine the impact of explant-derived factors on these distinct ovarian subcompartments (54, 55) (Fig. 4A). Organoid and follicle transcriptomic analyses were performed after 48 hr of treatment, while follicle growth, survival, and oocyte maturation were assessed using eIVFG.

Treatment with conditioned medium upregulated 142 and downregulated 80 genes in stromal organoids, whereas 267 genes were upregulated and 113 were downregulated in treated secondary follicles (Supplemental Fig. 4A,B). Only 22 upregulated genes were shared between the organoid and follicle models (Fig. 4C), indicating that factors released by MNGC-enriched explants induce largely compartment-specific responses. Pathway analysis further demonstrated that treatment with conditioned medium induced biologically distinct effects in each compartment. Top GO terms from genes upregulated in treated stromal organoids included “tissue remodeling,” “connective tissue development,” and “extracellular matrix organization” (Fig. 4D), consistent with the profibrotic remodeling programs that are observed in the aging ovarian stroma. In follicles, genes upregulated with treatment were enriched for inflammatory and immune-related pathways, including “inflammatory response,” “TNF signaling pathway,” and “cytokine-mediated signaling pathways,” among others (Fig. 4E), indicating acquisition of a proinflammatory state consistent with ovarian inflammaging. Together, these compartment-specific responses recapitulate key fibroinflammatory features of ovarian aging, with stromal remodeling occurring alongside follicular inflammation. Downregulated processes in organoids included multiple pathways related to solute transport, including “transport of small molecules” and “monatomic ion transmembrane transport”, indicative of suppressed transport and homeostatic programs in the stromal compartment (Supplemental Fig. 4C). In follicles, downregulated processes included “neuron projection development,” “cell morphogenesis,” and “signaling by BMP,” suggestive that processes such as transzonal projection formation and oocyte-granulosa cell communication may be disrupted (Supplemental Fig. 4D). Together, these findings demonstrate that factors secreted by MNGC-enriched explants are sufficient to induce distinct stromal-remodeling and follicular-inflammatory responses that parallel key fibroinflammatory phenotypes of the aging ovary. Thus, factors released by MNGCs and cells within their immediate microenvironment likely contribute to fibroinflammatory phenotypes that arise in the aging ovary.

To determine the functional consequences of sustained exposure to factors secreted by MNGC-enriched explants throughout folliculogenesis, explant-conditioned medium was utilized for long-term treatment of follicles in an eIVFG system that recapitulates folliculogenesis, oogenesis, and ovulation (54, 56). Treatment with conditioned medium transiently increased follicle growth relative to controls between day 2 (1.09-fold) and 8 (1.28-fold) with the greatest difference observed at D6 (1.35-fold) (Fig. 4F,G). However, follicle size did not differ between controls and treated follicles by the end of culture, and survival was comparable and >90% across culture between groups. Following ovulation, follicles that were treated with conditioned medium yielded a significantly lower proportion of mature metaphase II-arrested eggs (38.9%) compared to control follicles (82.3%) (Fig. 4H). These data indicate that secreted factors from MNGC-enriched explants transiently dysregulate follicle growth and impair oocyte maturation. Thus, the MNGC-enriched microenvironment is a source of signals that induce fibroinflammatory remodeling across ovarian compartments and compromise oocyte maturation, providing a functional link between MNGCs and key ovarian aging phenotypes.

## DISCUSSION

This study establishes failed post-ovulatory debris clearance as a previously unrecognized feature of ovarian aging that promotes fibroinflammaging. Using a novel CL-tracking strategy, we found that CL regression becomes dysregulated with advanced reproductive age, resulting in the accumulation of persistent CLs. These persistent CLs represent a unique debris burden to the ovary, as they are fibroinflammatory foci characterized by collagen and lipid accumulation, immune cell enrichment, and an inflammatory transcriptomic signature. pCLs also exhibit high transcriptomic similarity to ovarian MNGCs and *ex vivo*-derived CLs can acquire MNGC-like characteristics, suggesting that pCLs are closely associated with and may promote MNGC formation in the aging ovary. Finally, we demonstrated that factors secreted by MNGCs and their associated microenvironment induce gene signatures consistent with stromal remodeling and follicle inflammation, and that long-term exposure of follicles results in impaired oocyte development. Together these findings support a model in which dysregulated debris clearance results in the accumulation of fibroinflammatory pCLs, which promote MNGC formation that in turn mediate fibroinflammation of multiple ovarian compartments (Fig.5). Thus, dysregulated debris clearance and its downstream consequences are likely major contributors to ovarian fibroinflammaging.

**Figure 5.**
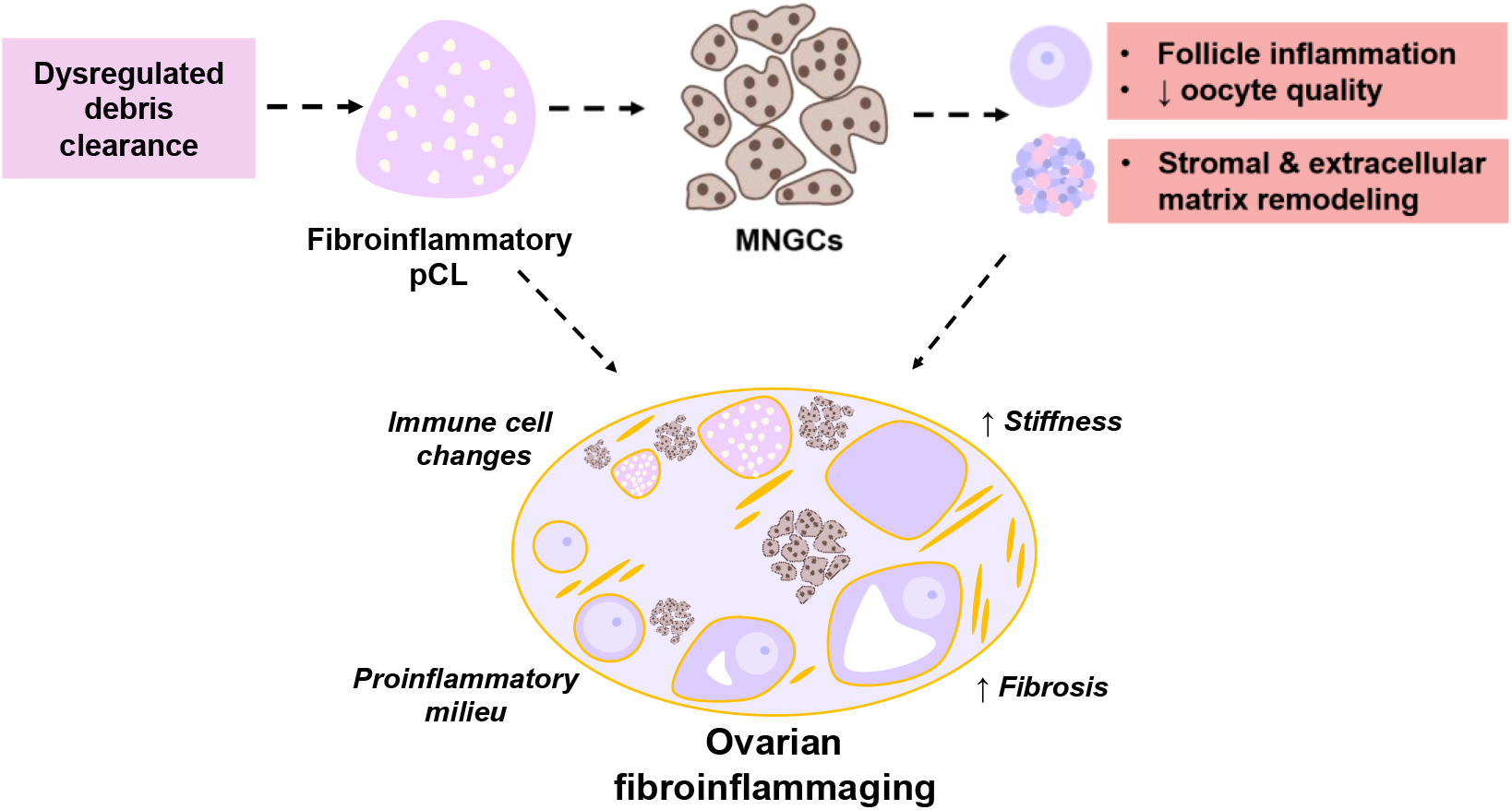
Graphical Summary. Schematic overview demonstrating the contribution of age-associated dysregulated debris clearance and its downstream consequences to ovarian fibroinflammaging.

Our results of dysregulated CL regression and the resulting accumulation of pCLs support the idea that the cumulative burden of repeated ovarian remodeling contributes to loss of tissue homeostasis with age. Consistent with this, mouse models in which ovulatory burden are reduced, such as through continuous breeding, exhibit decreased age-associated ovarian fibrosis (57). Additionally, in humans, conditions such as parity, breastfeeding, and oral contraceptive use are associated with delayed menopause and decreased risk of early menopause, suggesting that reduced ovarian cyclicity and ovulation may extend reproductive function (58, 59). Thus, repeated cycles of ovarian remodeling may place a substantial demand on debris clearance mechanisms over the reproductive lifespan, such that debris accumulation eventually outpaces clearance as these processes decline with age. Importantly, CL regression is only one source of ovarian debris. Follicular atresia as well as ovulation-associated tissue damage and subsequent wound repair also generate substantial cellular debris that must be continually cleared and may similarly contribute to age-associated loss of ovarian homeostasis. In fact, abrogated atretic follicle debris clearance in IL-33 knockout mice results in accelerated MNGC development, reduced autophagic activity, and shortened reproductive lifespan, further linking impaired debris clearance with ovarian aging (60). Thus, failed post-ovulatory debris clearance is one contributor to a broader multi-factorial age-associated loss of ovarian tissue homeostasis.

A central finding of this study is the close molecular relationship between pCLs and ovarian MNGCs. pCLs express both CL- and MNGC-specific markers, identifying them as putative intermediates, and this phenotype is recapitulated in an *ex vivo* model of CL persistence. pCLs also likely contain macrophages and T cells in addition to luteal cells and are enriched for collagen and lipids, features which may promote the acquisition of an MNGC phenotype by macrophages. Macrophages can fuse when they encounter difficulty engulfing material, a process termed “frustrated phagocytosis,” which can trigger an inflammatory response (61–63). Consistent with this, *Gpnmb* is highly expressed in ovarian and non-ovarian MNGCs, including osteoclasts, foreign body giant cells, and Langhans giant cells, as well as lipid-associated macrophages associated with obesity, atherosclerosis, and neurodegenerative diseases such as Alzheimer’s Disease, Parkinson’s Disease, and Multiple Sclerosis (45, 49, 50, 64–70). Thus, *Gpnmb* expression in pCLs and persistent *ex vivo* CLs may reflect activation of a debris-processing response associated with lipid accumulation, phagocytic stress, and acquisition of MNGC-associated phenotype. However, the cellular origin of these signatures remains unclear, as our LCM-based bulk RNAseq approach cannot resolve cell-specific contributions within the heterogeneous CL. The intermediate pCL profile may therefore reflect macrophages that infiltrate the persistent luteal environment and acquire MNGC-like properties. Alternatively, the acquisition of *Gpnmb* in the *ex vivo* CL model raises the possibility that luteal cells themselves may acquire some MNGC-associated characteristics. This is consistent with the known plasticity, phagocytic capacity, and immune signaling functions of granulosa cells (71–73), although whether these properties are retained after luteinization remains unknown. Because our BrdU-tracking experiment ended at 3 wks post-ovulation, it likely did not extend sufficiently to capture later MNGC formation at persistent CL sites. Extending this time course, together with lineage-tracing and single-cell analyses, will help resolve the cellular populations that contribute to the pCL microenvironment and directly give rise to ovarian MNGCs.

MNGC-enriched factors were found to modulate fibroinflammatory gene expression in an ovarian compartment-specific manner. Factors released by MNGC-enriched explants induced signatures associated with tissue and extracellular matrix remodeling in stromal organoids, features consistent with age-associated remodeling of the ovarian stroma in vivo (1, 2, 5, 55). The MNGC-enriched secretome also promoted inflammatory signaling in follicles, with associated dysregulated folliculogenesis and a decrease in the proportion of oocytes that reached metaphase II and capable of undergoing fertilization. These effects parallel the age-associated inflammatory signature and the established decline in follicle quality with age (9, 74). This physiological impact on folliculogenesis, a process essential to both fertility and steroid hormone production, indicates that MNGCs and their immediate microenvironment have the potential to negatively impact reproductive longevity. Thus, while MNGCs likely arise in response to the accumulation of pCL debris, they can also negatively impact ovarian function and contribute to ovarian fibroinflammaging. Together, these findings suggest that failed debris clearance and the resulting accumulation of pCLs and MNGCs may contribute to a self-perpetuating fibroinflammatory environment within the aging ovary.

The identification of dysregulated CL regression, persistent luteal debris accumulation, and MNGC formation as contributors to ovarian inflammaging highlights several potential points of therapeutic intervention to support ovarian longevity. For example, restoration of effective CL regression in aged ovaries could prevent pCL accumulation through methods that enhance autophagy, efferocytosis, or other age-affected debris clearance processes. Additional characterization of these cellular processes within the ovary and to age-associated debris accumulation will aid in determining which mechanisms represent promising therapeutic targets to mitigate debris-induced fibroinflammation. Additionally, the distinctive molecular features of pCL debris and MNGCs may make these structures uniquely targetable. For instance, molecular markers such as GPNMB, could enable selective targeting of both pCLs and MNGCs within the ovary. In somatic tissues, GPNMB is upregulated in various pathologies, and efforts are underway to target GPNMB therapeutically in the context of various cancers and senescence (75–78). Recently, a GPNMB vaccine developed using GPNMB-derived peptides successfully reduced senescent cells, ameliorated aging phenotypes, and extended lifespan in a progeroid mouse model (78). Thus, GPNMB may be a candidate therapeutic target to mitigate the downstream fibroinflammatory consequences of debris accumulation. Overall, our findings demonstrate the potential for targeting the upstream drivers and downstream consequences of ovarian debris to mitigate ovarian inflammaging.

In summary, this study identified failed post-ovulatory debris clearance as a previously unrecognized contributor to ovarian fibroinflammaging. We have established a relationship between debris clearance failure, pCL accumulation, and MNGC formation in the aged ovary, with each sequential event contributing to a fibroinflammatory environment. These findings connect dysregulation of homeostatic cellular and tissue maintenance processes with multiple hallmarks of ovarian aging and identify debris clearance as a promising therapeutic target to modulate ovarian inflammaging and extend ovarian functional longevity.

## MATERIALS AND METHODS

### Animals

All experiments were performed using C57Bl/6J and C57Bl/6Hsd mice purchased from Jackson Laboratories (Bar Harbor, ME, USA) or Envigo (Indianapolis, IN, USA), respectively. Reproductively young mice between 6-12 wks and reproductively old mice between 9-18 months were used for all analyses unless otherwise specified. Upon arrival to Northwestern University, all mice were housed in a controlled barrier facility at the Center for Comparative Medicine (Chicago, IL, USA). Mice were housed under constant light (14-hour light/10-hour dark), temperature, and humidity control. Mice received water and chow ad libitum. All animal use and procedures were approved by Northwestern University’s Institutional Animal Care and Use Committee.

### Tissue processing, histochemical staining, and imaging

Ovaries were fixed in Modified Davidson’s Solution (Electron Microscopy Sciences, Hatfield, PA, USA) overnight, washed in 70% ethanol, and stored at 4°C until processing. Samples were dehydrated and processed using an automated tissue processor (Leica Biosystems, Buffalo Grove, IL, USA) and then embedded in paraffin and sectioned at a thickness of 5 µm using a Reichert-Jung Biocut 2035 microtome (Leica Biosystems). Hematoxylin and Eosin (H&E) staining of every 5^th^ slide was conducted using a Leica Autostainer XL (Leica Biosystems). Samples were mounted with Cytoseal XYL (Thermo Fisher Scientific, Waltham, MA, USA). Imaging was performed with an EVOS FL Auto Cell Imaging System (Thermo Fisher Scientific) using 20x and 40x objectives. Tiled images were stitched utilizing EVOS imaging software to achieve images of full ovarian sections.

### Ovarian superovulation and corpora lutea tagging

Reproductively young and old mice were administered 5 IU pregnant mare serum gonadotropin (PMSG) (ProSpec, East Brunswick, NJ, USA) via intraperitoneal (IP) injection to induce ovarian hyperstimulation. After 44-46 hrs, mice were administered 5 IU human chorionic gonadotropin (hCG) (Sigma-Aldrich, St. Louis, MO, USA) via IP injection to induce ovulation. Because ovulation kinetics are delayed with age (8, 79), the initial bromodeoxyuridine (BrdU; 100µg/g body weight; Abcam, Waltham, MA) IP injection was performed at 13 h and 16 h following the hCG ovulatory trigger in reproductively young and old mice, respectively. To ensure successful tagging across the entire ovulatory event, a second BrdU injection was performed 12 h following the initial injection when CL angiogenesis is at its maximum (80). Mouse ovaries were sampled 1 day, 1 wk, 2 wks, and 3 wks post-ovulation.

### Immunohistochemistry

Ovarian samples were deparaffinized and rehydrated using Citrosolv and descending ethanol concentrations. Antigen retrieval was performed with Reveal Decloaker (Biocare Medical, Pacheco, CA, USA) prior to blocking with the Avidin/Biotin Blocking kit (Vector Laboratories, Newark, CA, USA) and 10% normal goat serum (Vector Laboratories) according to the manufacturer’s instructions. Samples were incubated with primary antibodies for BrdU (1:500; Abcam; Cat#: ab6326), CD3ε (1:400; Cell Signaling Technology; Cat#: 78588), or F4/80 (1:50; Bio-Rad, Hercules, CA, USA; Cat#: MCA497) overnight at 4°C. Samples were washed in Tris-buffered saline with 0.1% Tween-20 (Sigma Aldrich) (TBS-T) and incubated with a biotinylated anti-rat or anti-rabbit secondary antibody (1:200, BA-9401 and PK-6101; Vector Laboratories) for 1 hour at room temperature. Signal amplification was performed with a Vectastain Elite ABC Kit (Vector Laboratories) and detection was performed with a 3,3′-diaminobenzidine (DAB) using a DAB Peroxidase (HRP) Substrate Kit (Vector Laboratories). Samples were counterstained with hematoxylin, then dehydrated with ascending ethanol concentrations and Citrosolv and mounted with Cytoseal XYL (Epredia, Kalamazoo, MI, USA). Sections were imaged with an EVOS M7000 (Thermo Fisher Scientific) at 20X and image analysis was performed using Fiji (NIH). Color deconvolution was performed to separate the hematoxylin from DAB staining. For immune cell area quantification, percent positive CD3ε or F4/80 area relative to total area of a region of interest (ROI) that included area of a CL or pCL was determined. Up to 5 CLs or pCLs from a single ovarian section were quantified for a minimum of 3 animals.

### Autophagy analysis

Reproductively young (4 m) and old (10-11 m) mice were stimulated to undergo superovulation, and CLs were microdissected from ovaries 7 days post hCG injection. CLs were pooled from individual mice and homogenized in 100 µl RIPA buffer (Thermo Fisher Scientific) containing protease inhibitor (Thermo Fisher Scientific) by vortexing for 1 hr at 4°C. The protein lysates were centrifuged at 10,000 g for 3 min to pellet insoluble material. Laemmli sample buffer (Bio-Rad) and β-mercaptoethanol (10%; Sigma-Aldrich) were added to each sample, which were then boiled for 10 min and stored at -20°C until analysis by Western blot. Electrophoresis was performed on 10 µg of CL protein loaded on 4-15% premade SDS-polyacrylamide gels (Bio-Rad). Resolved proteins were transferred to PVDF membranes, blocked with 5% bovine serum albumin (BSA; Sigma-Aldrich) for 2 h, and incubated in primary antibodies (LC3A/B, 1:1,000; Cell Signaling Technologies (Cat#12741), Danvers, MA, USA) diluted in in 5% BSA overnight at 4°C. Membranes were washed in TBS-T, incubated in HRP-conjugated secondary antibody (1:10,000, Cytiva, Wilmington, DE, USA) for 1 h at room temperature, washed, and protein-antibody complexes were detected using Amersham ECL Plus chemiluminescence reagents (Cytiva). Blots were then stripped using Restore Western Blot Stripping Buffer (Thermo Fisher Scientific), re-blocked, and probed with antibodies against α-tubulin for normalization of protein loading. Protein bands were visualized using a ChemiDoc (Bio-Rad) and band densitometry was assessed by Fiji.

### Persistent corpus luteum (pCL) and multinucleated giant cell quantification

pCLs were identified by their distinguishing morphological phenotype and were quantified by assessing every 15^th^ section of each ovary, avoiding duplicate counting of the same pCLs. Quantification was performed on three ovaries from reproductively young and old mice (12 m). To determine MNGC penetrance in the ovary, the Trainable Weka Segmentation plugin in Fiji (NIH) was trained to identify non-ovary, non-MNGC ovary, and MNGC regions as previously described (45). MNGC quantification was performed on three sections from different ovarian regions, with the average MNGC area being reported.

### Laser capture microdissection

CLs, pCLs, and MNGCs were collected using Zeiss Palm MicroBeam Laser Capture Microdissection system (Carl Zeiss Microscopy, Jena, Germany). Fresh frozen samples embedded in optimal cutting temperature (OCT) compound were cryosectioned at 10 µm thickness under RNase-free conditions, adhered to nuclease-free PEN membrane slides (Carl Zeiss Microscopy), and stored at -80°C. Serial sections were placed on glass slides for subsequent H&E staining to confirm the presence of structures of interest. Immediately prior to laser capture microdissection (LCM), tissue slides were fixed in 70% RNase-free ethanol (Thermo Fisher Scientific), transferred into RNase-free water (Corning, Manassas, VA, USA) to remove OCT, then dehydrated in 70% RNase-free ethanol followed by three 100% RNase free ethanol washes (3 min each), and then air dried. The Zeiss Palm was calibrated for cut energy, cut focus, laser pressure catapulting (LPC) energy, and LPC focus on the 20x objective. CLs, pCLs, and MNGCs were identified by their distinct morphological characteristics. All cuts were outlined using the RoboLPC setting and free-hand tool for selective capture of tissue structures. Tissue was collected in the Zeiss AdhesiveCap 500 clear microtube (Carl Zeiss Microscopy, Göttingen, Germany). A minimum area of 250,000 µm^2^ of tissue was collected for each sample, and different biological pools were obtained from 4 mice. Samples were stored in a -80°C until RNA isolation.

### RNA isolation and bulk RNA sequencing

A RNeasy Micro Kit (Qiagen, Germantown, MD) was used to isolate total RNA from LCM samples, per the manufacturer’s instruction. 350 µl of RLT buffer containing β-mercaptoethanol was added to the AdhesiveCap microtube, vortexed for 30 s, and incubated upside down for 30 min at room temperature prior to moving samples to 1.5 ml RNase-free centrifuge tubes for RNA isolation. RNA integrity was determined using a 2100 Bioanalyzer System (Agilent Technologies, Inc., Santa Clara, CA, USA) with a Bioanalyzer RNA Pico chip (Agilent Technologies). All LCM-collected RNA samples had an RIN of 6.8 or greater.

The SMART-Seq v4 Ultra Low Input RNA Kit (Takara Bio, San Jose, CA, USA) was used to generate cDNA according to the manufacturer’s protocol. The Qubit DNA HS assay and Agilent Bioanalyzer DNA HS chip were used to measure resulting cDNA quantity and quality, respectively. High quality cDNA (greater than 800 bp) was used to generate sequencing libraries. The Nextera XT DNA library preparation kit (Illumina, Inc., San Diego, CA, USA) was used for library preparation, using 200 ng of cDNA. Multiplexed libraries were sequenced on the Novaseq X Plus at Northwestern University’s NUseq Core. FastQC was used to assess the quality of the reads in FASTQ format, and reads were trimmed from the 3’ ends using cutadapt to remove Illumina adapters. Using STAR, the trimmed reads were aligned to the *Mus musculus* genome (mm10) (81). Read counts were calculated using a gene annotation file from Ensembl (http://useast.ensembl.org/index.html) and htseq-count (82). Data normalization and differential expression analyses were performed with DESeq2 and the contrast argument of the results function was utilized to compare structures to one another or between control and treatment groups (83). Genes were considered significantly differentially expressed (DEGs) between groups if they had an adjusted *P*-value < 0.05. Data were log transformed and principal component analysis plots were generated using the plotPCA function. Correlation plots were made in SRplot using the normalized count matrix. Volcano plots were created using the EnhancedVolcano package in R (version 1.20.0). Gene Ontology analyses were performed using Metascape, prioritizing DEGs with an absolute log_2_ fold-change ≥1 (ovarian-specific structures) or ≥ 0.5 (follicle and organoids). All heatmaps were created using SRPlot.

### In situ RNA hybridization

RNAscope kits were used to visualize mRNA transcripts of *Ptgfr* and *Gpnmb* (Advanced Cell Diagnostics, Newark, CA, USA) in young and old mouse ovarian sections as well as sectioned *ex vivo* CLs, per the manufacturer’s instructions. Imaging analysis was performed using Fiji. Color deconvolution was performed to separate the hematoxylin and RNAscope probe positive signal.

### Picrosirius red staining

Ovarian tissue sections were deparaffinized and rehydrated using Citrosolv and descending ethanol concentrations. Tissue sections were then incubated in picrosirius red (PSR) staining solution, prepared following standard protocol (1), for 40 min. Tissue sections were then incubated in an acidified water solution (0.5M hydrochloric acid; Thermo Fisher Scientific) for 90 seconds. Samples were dehydrated in three subsequent 100% ethanol washes and cleared using Citrosolv for 5 min. Slides were mounted using Cytoseal XYL (Epredia) then imaged. Bright-field images were taken using the EVOS FL Auto Cell Imaging System (Thermo Fisher Scientific) using 20x and 40x objectives. ImageJ was used to quantify positive PSR staining above a set threshold which was kept constant for all samples. ROIs were defined for each CL or pCL per ovary and 1-5 CLs or pCLs from a single ovarian section were quantified for a minimum of 3 animals.

### Oil Red O Staining

Oil Red O staining was performed per the manufacturer’s instructions (Abcam) on frozen ovarian tissue sections that were fixed with 4% paraformaldehyde for 10 minutes prior to staining. Tissue samples were counterstained with Hematoxylin, rinsed with water, then mounted using an aqueous mounting medium.

### Ex vivo CL regression assay

Multilayer secondary follicles were isolated through enzymatic digestion of prepubertal (postnatal day 12-16) ovaries as previously described (54). Follicles were encapsulated in 0.5% alginate and cultured at 37°C in a humidified atmosphere of 5% CO_2_. On day 8 of culture, follicles were removed from alginate by incubation with 10 IU/ml alginate lyase and transferred to maturation medium (αMEM-Glutamax containing 10% fetal bovine serum, 10 ng/ml mouse epidermal growth factor (EGF; BD Biosciences, Franklin Lakes, NJ, USA), 1.5 IU/ml hCG (Sigma-Aldrich) and 10 mIU/ml Gonal-F) for 16 h at 37°C to induce ovulation and luteinization. Oocytes were removed and post-ovulatory follicles were transferred to αMEM containing 10% FBS and 1% penicillin-streptomycin (PS) in ultra-low adherent 96-well plates for CL culture. Half-volume medium changes were performed every other day, with subsets of CLs being fixed and processed for histological analyses on 2 days, 1 wk, and 2 wks post-ovulation.

### Generation of MNGC-enriched explant conditioned medium

Intact reproductively old ovaries (12-15 m) were placed in digestion medium containing Leibowitz L15 medium with 1% fetal bovine serum (FBS; Gibco, Waltham, MA), 0.5% PS, 0.4 mg/ml Collagenase IV (205 units/mg; Gibco), and 0.2 mg/ml DNase I (Sigma Aldrich) for 30 min on a heated bench. During digestion, ovarian tissue and structures such as CLs and large follicles were resected to expose the stromal-enriched interior. At this point, MNGCs were visualized by their pigmented appearance by brightfield microscopy (Fig. 4Bi) and microdissected. After the 30 min enzymatic digestion, dissection of MNGC-enriched fractions was continued in L15 medium without enzymes. Non-pigmented tissue was resected from putative MNGC-containing fragments, and the fractions were confirmed to be autofluorescent using a FL EVOS Auto Imaging System (Thermo Fisher Scientific) (Fig. 4Bii). Individual MNGC-enriched explants were placed into 100 µl DMEM containing 10% FBS + 1% PS in an ultra-low attachment 96-well plate overnight. Autofluorescence was confirmed again the next day, with any explants that no longer exhibited autofluorescence being discarded. MNGC-enriched explants were then transferred to either follicle base medium (αMEM-Glutamax medium containing 3 mg/ml bovine serum albumin (BSA) (MP Biomedicals, Santa Ana, CA, USA) and 1 mg/ml fetuin (Sigma-Aldrich)) or organoid treatment medium (Dulbecco’s Modified Eagle Medium (DMEM) containing 1% PS) and incubated for 24 hrs. After conditioning, conditioned medium was collected and frozen for up to 2 wks at -20°C. Medium not exposed to MNGC-enriched explants was frozen and used as the control treatment.

### Proinflammatory cytokine analysis

MNGC-enriched explant conditioned medium samples (conditioned for 24 or 48 hr) were thawed and analyzed using a Quantibody Mouse Inflammation Array (RayBiotech, Peachtree, GA, USA; Cat#: QAM-INF-1-1) following the manufacturer’s protocol. Conditioned medium samples were diluted 2-fold with sample diluent prior to use. Non-conditioned medium controls were used as negative controls. Array slides were scanned and analyzed by RayBiotech.

### Treatment of ovarian stromal organoid and secondary follicles with MNGC-enriched explant conditioned medium

Reproductively young mice were used to isolate and generate early secondary follicles and ovarian stromal organoids as previously described (54, 55). Organoids were allowed to aggregate for 48 hr prior to treatment, whereas treatment of follicles was started on the day of isolation. MNGC-enriched explant conditioned medium was diluted 1:1 with αMEM-Glutamax medium containing 3 mg/ml BSA, 1 mg/ml fetuin, 0.2% insulin-transferrin-selenium (Thermo Fisher Scientific), and 20 mIU/ml Gonal-F for subsequent follicle treatment. For organoid treatment, conditioned medium was diluted 1:1 in fresh DMEM. For control treatments, frozen control medium was diluted 1:1 as described above. Follicles from the same biological pool and organoids generated within the same micromold were split between control and conditioned-medium groups. Treatments were performed for 48 hrs after which follicles and organoids were snap frozen in RLT buffer containing β-mercaptoethanol. RNA isolation and bulk RNA sequencing were carried out as described above. For long-term treatment of follicles with conditioned medium, fresh aliquots of conditioned medium were thawed and used for 50% medium changes every other day of culture. Follicle survival, growth, and meiotic maturation resumption were assessed as previously described (54, 84).

### Statistical analysis

Statistical analyses were performed using GraphPad Prism software (10.2.3) or R (RStudio, version 4.3.0). Statistical tests were selected according to the experimental design and data type. All data were tested for normality, and appropriate statistical tests were selected based on data distribution. For comparisons among three or more groups, data were analyzed using either one-way ANOVA with post-hoc Tukey’s multiple comparison test (normal distribution) or Kruskal-Wallis test with Dunn’s multiple comparison test. For comparison between two groups Welch’s unpaired *t* test was used. Categorical outcomes were analyzed using a χ2 test. Data are represented as Mean ± SEM. *P < 0.05* was considered significant. Specific statistical tests and sample size are indicated in the corresponding figure legends.

## DATA, MATERIALS, AND SOFTWARE AVAILABILITY

Raw data and expression count matrices are deposited in GEO (Accession numbers GSE347815 (laser capture dissected structures) and GSE348355 (MNGC-enriched conditioned medium modulation of follicles and organoids). Differential gene expression analysis codes have been uploaded to Northwestern University Galter Health Sciences Library Prism https://doi.org/10.18131/qafrc-pv225.

## ACKNOWLEDGMENTS

We acknowledge the Northwestern University’s Center for Advanced Microscopy (CAM) and NUseq Core for support with laser capture microdissection training and bulk RNAseq experiments, respectively. We acknowledge Desiree Franklin for her assistance in optimizing the ex vivo corpus luteum culture system.

---

**Supplemental Figure 1.**
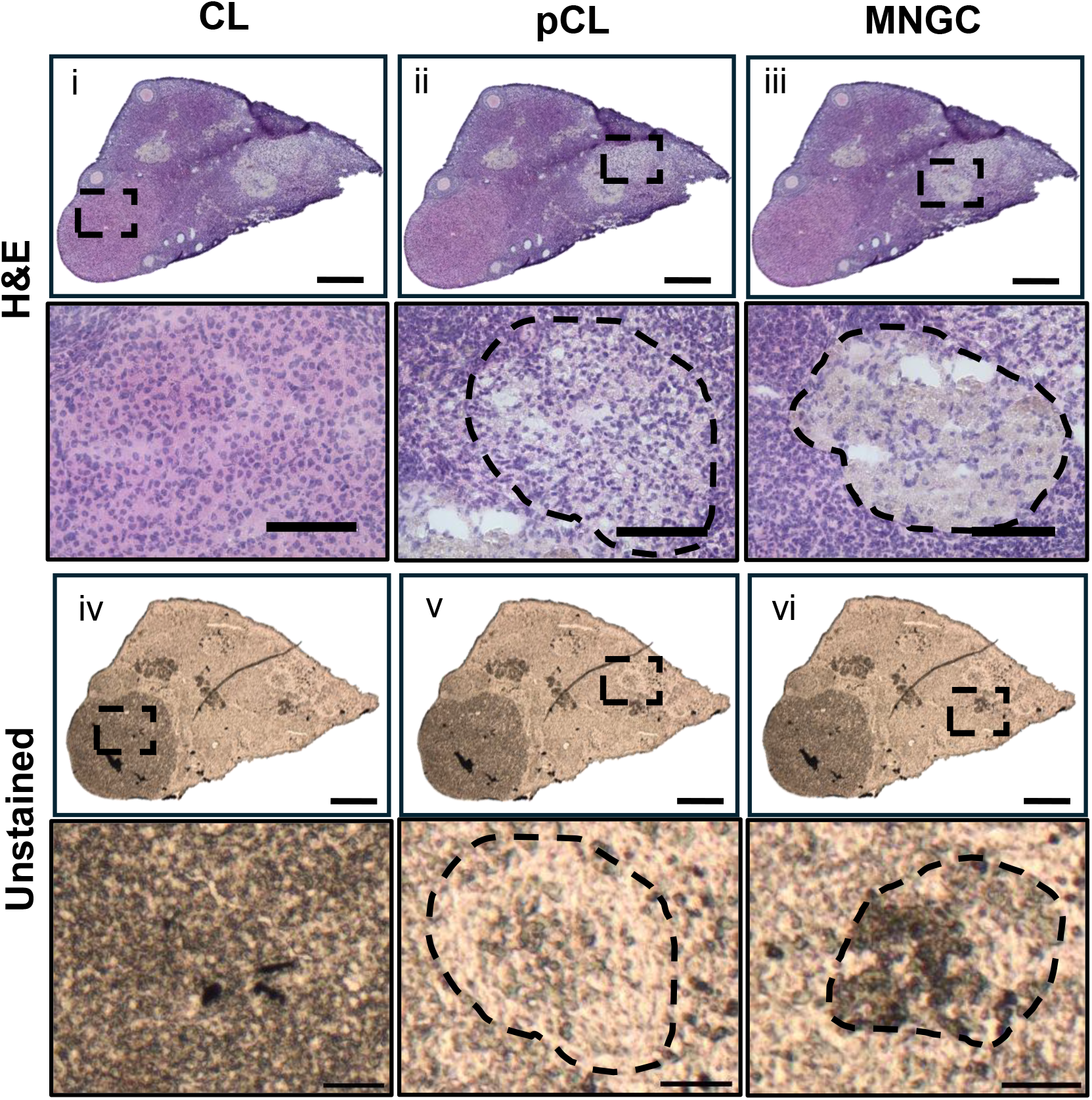
Identification of CL, pCL, and MNGC samples for laser capture microdissection. Representative images of unstained ovarian tissue sections (i-iii) and representative H&E-stained serial sections (iv-vi) showing morphology of CLs (i, iv), pCLs (ii, v), and MNGCs (iii, vi). Scale bars: Top panes (whole ovary scans) 300 µm; lower panels (magnified images of CLs, pCLs, MNGCs) 100 µm.

**Supplemental Figure 2.**
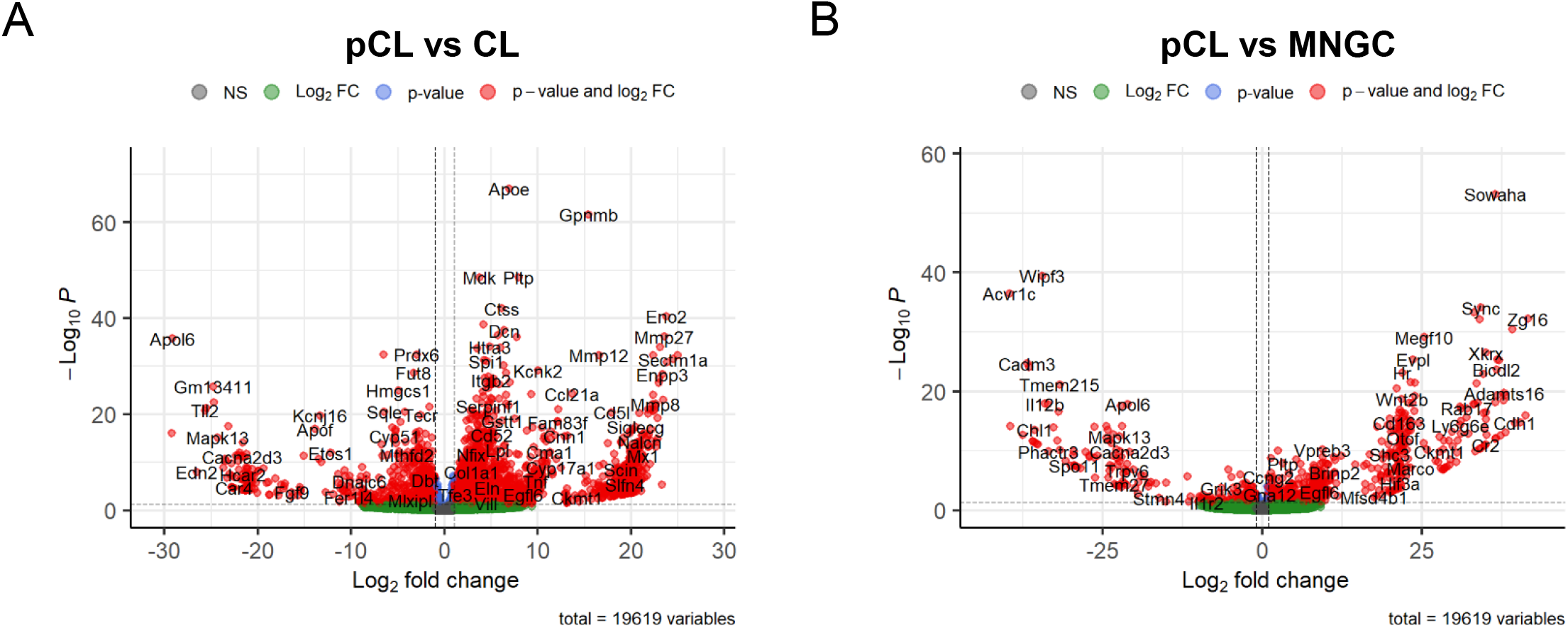
Differential gene expression analysis of pCLs. (A) Volcano plot of differentially expressed genes (DEGs) between pCLs and CLs. (B) Volcano plot of DEGs between pCLs and MNGCs. Genes notated in red indicate adjusted *P* < 0.05 and Log_2_FC ≥ 1.

**Supplemental Figure 3.**
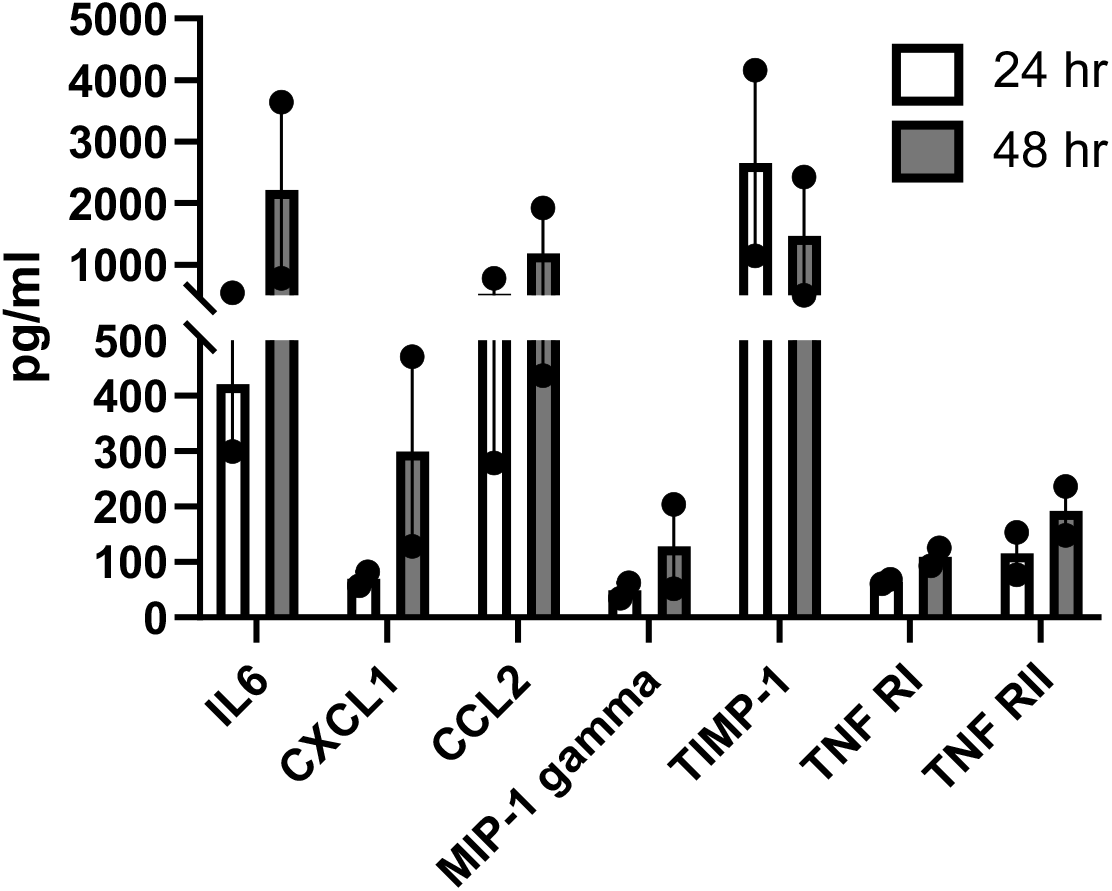
Proinflammatory mouse cytokine analysis of MNGC-enriched explant conditioned media. Media was conditioned for 24 or 48 hr. 2 biological samples from pooled MNGC-enriched explant conditioned media were assessed for each timepoint.

**Supplemental Figure 4.**
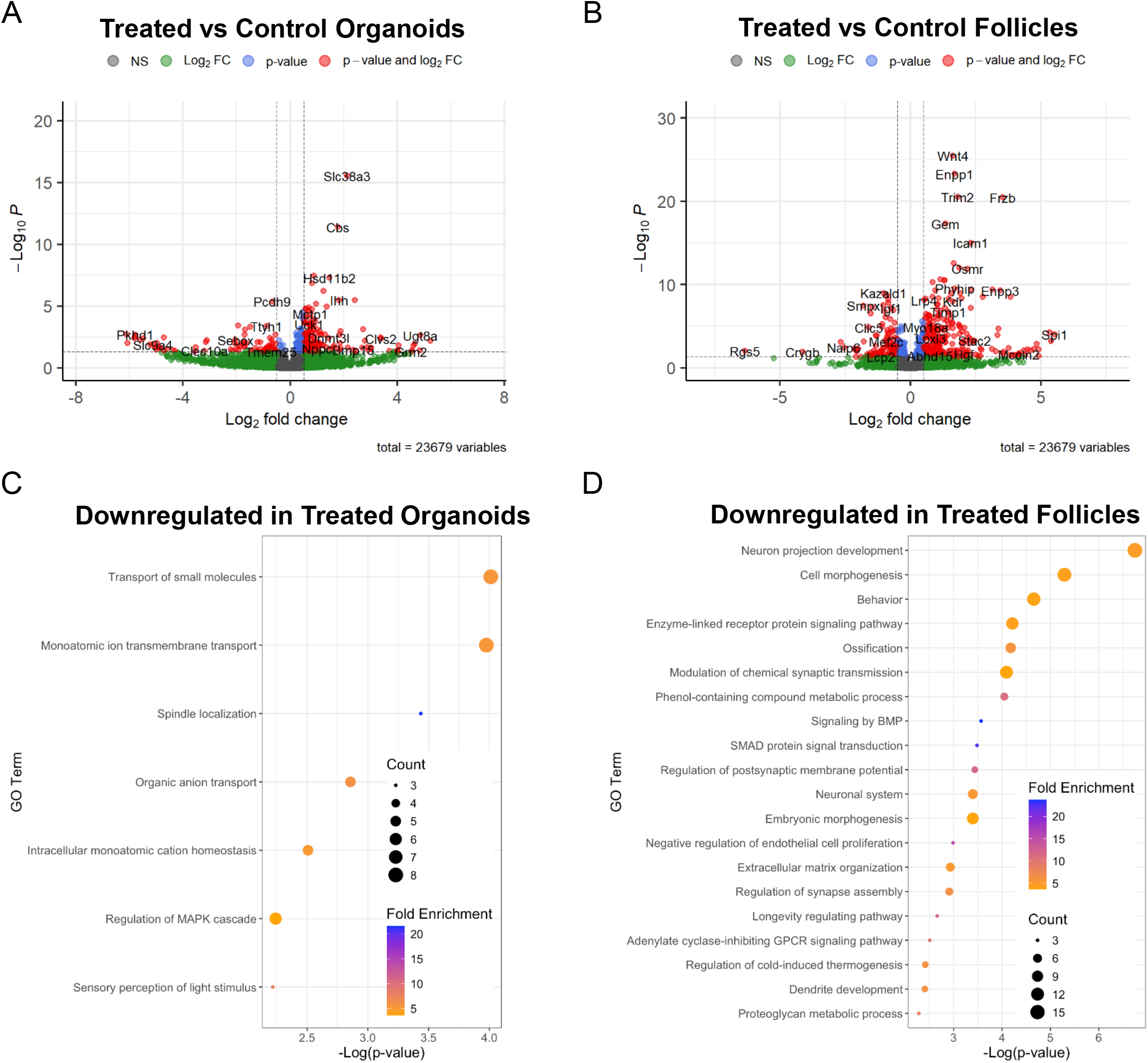
Transcriptomic analysis of MNGC-enriched explant conditioned media-treated follicles and stromal organoids. Volcano plots of DEGs between conditioned media-treated and control ovarian stromal organoids (A) and follicles (B). Genes notated in red indicate adjusted *P* < 0.05 and Log_2_FC ≥ 0.5. Pathway analysis of downregulated DEGs (conditioned media-treated vs control) from stromal organoids (C) and follicles (D).

