## Supplemental Figures for "Age-dependent dysregulated post-ovulatory debris clearance promotes ovarian fibroinflammation"

**Supplemental data for**
**Age-dependent dysregulated post-ovulatory debris**
**clearance**
**promotes ovarian fibroinflammation**

Aubrey Converse<sup>1,†,\*</sup>, Madeline J. Perry<sup>1,†</sup>, Shweta S. Dipali<sup>1</sup>,
Prianka H. Hashim<sup>1</sup>, Michele T. Pritchard<sup>2</sup>, Francesca E.
Duncan<sup>1,\*</sup>

<sup>1</sup> Department of Obstetrics and Gynecology, Feinberg School of
Medicine, Northwestern University, Chicago IL 60611, USA; <sup>2</sup>
Department of Pharmacology, Toxicology, & Therapeutics,
Institute for Reproductive and Developmental Sciences,
University of Kansas Medical Center, Kansas City, KS 66160,
USA

<sup>†</sup>Equal contributors

<sup>\*</sup>Co-corresponding authors

### Supplemental Figure 1

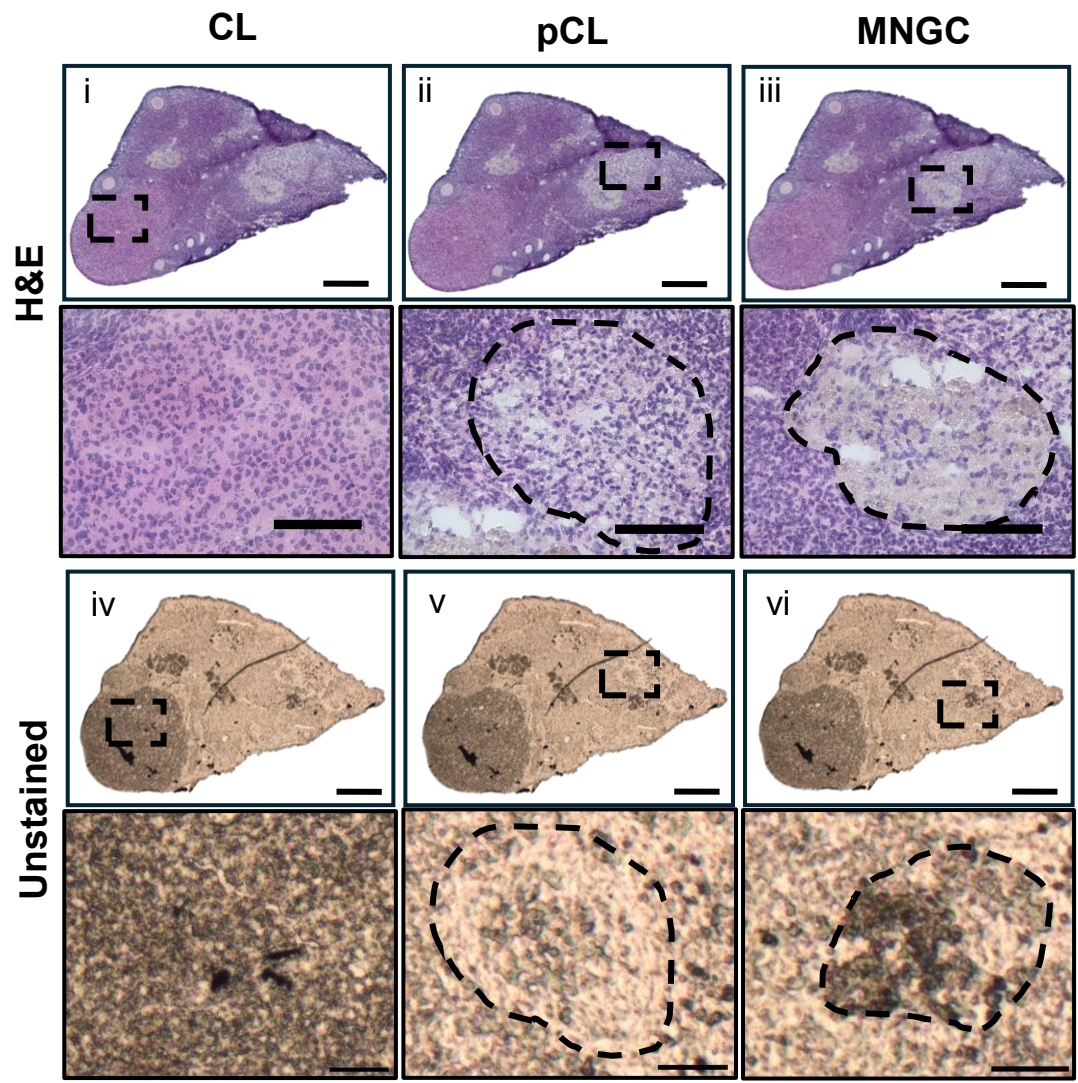

Supplemental Figure 1. Identification of CL, pCL, and MNGC samples for laser capture microdissection. Representative images of unstained ovarian tissue sections (i-iii) and representative H&E-stained serial sections (iv-vi) showing morphology of CLs (i, iv), pCLs (ii, v), and MNGCs (iii, vi). Scale bars: Top panes (whole ovary scans) 300 μm; lower panels (magnified images of CLs, pCLs, MNGCs) 100 μm.

A

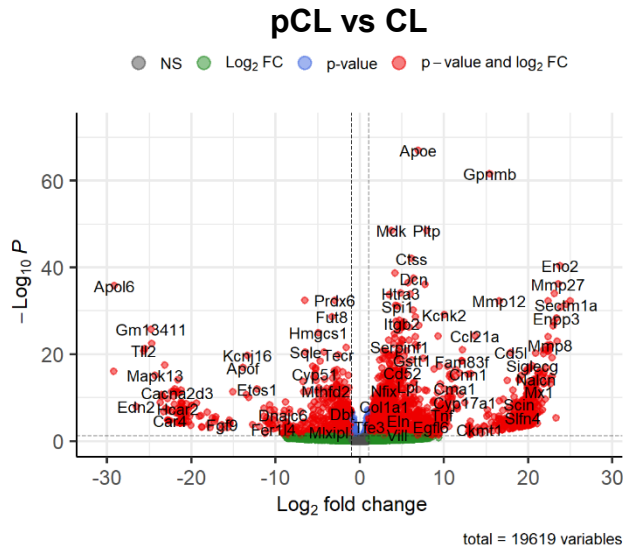

total = 19619 variables

# B

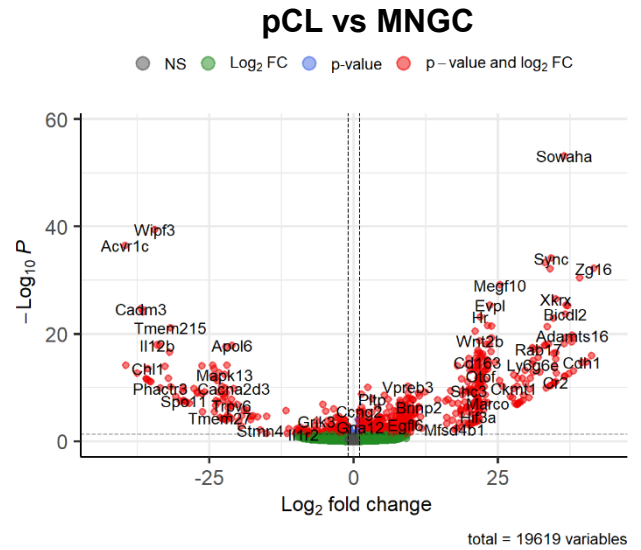

total = 19619 variables

Supplemental Figure 2. Differential gene expression analysis of pCLs. (A) Volcano plot of differentially expressed genes (DEGs) between pCLs and CLs. (B) Volcano plot of DEGs between pCLs and MNGCs. Genes notated in red indicate adjusted $P < 0.05$ and $\text{Log}_2\text{FC} \geq 1$.

### Supplemental Figure 3

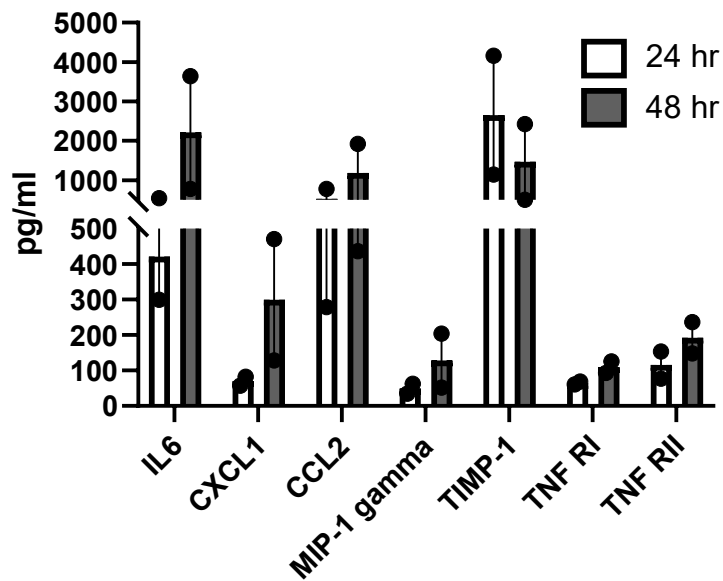

Supplemental Figure 3. Proinflammatory mouse cytokine analysis of MNGC-enriched explant conditioned media. Media was conditioned for 24 or 48 hr. 2 biological samples from pooled MNGC-enriched explant conditioned media were assessed for each timepoint.

### Supplemental Figure 4

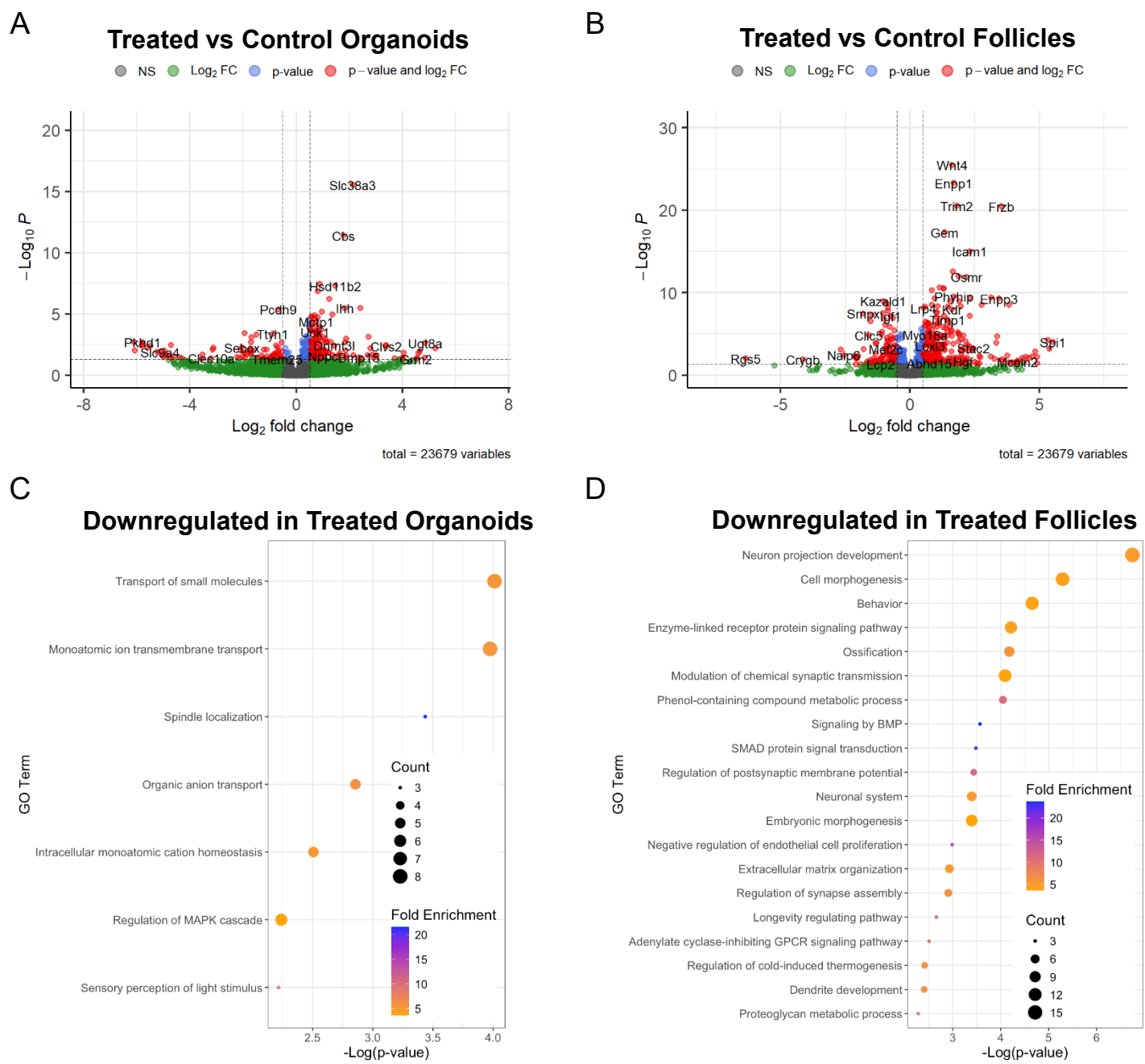

Supplemental Figure 4. Transcriptomic analysis of MNGC-enriched explant conditioned media-treated follicles and stromal organoids. Volcano plots of DEGs between conditioned media-treated and control ovarian stromal organoids (A) and follicles (B). Genes notated in red indicate adjusted $P < 0.05$ and $\text{Log}_2\text{FC} \geq 0.5$. Pathway analysis of downregulated DEGs (conditioned media-treated vs control) from stromal organoids (C) and follicles (D).
